# Novel characterization of peptidergic release from prefrontal cortical somatostatin neurons

**DOI:** 10.1101/599829

**Authors:** Nigel Dao, Dakota F. Brockway, Nicole A. Crowley

**Affiliations:** Department of Biobehavioral Health, Pennsylvania State University, University Park PA 16802; Neuroscience Curriculum, Pennsylvania State University, University Park PA 16802

## Abstract

Somatostatin is a neuropeptide thought to play a role in a variety of neuropsychiatric disorders, and is important for healthy aging and behavioral resiliency. Conditions governing somatostatin peptidergic release are not well-defined. Using a combination of optogenetic, transgenic, and biochemical approaches, we demonstrate an assay for the induction and inhibition of somatostatin in acute brain slices.

---

Somatostatin (SST) is a neuropeptide found throughout the brain^1^. SST-positive (SST+) neurons have begun to receive attention for their role in a wide variety of neuropsychiatric conditions and brain cancer^2^. Reduced SST has been found in the cerebrospinal fluid (CSF) of patients with major depressive disorder (MDD)^3^, schizophrenia and dementia^4^, and SST is thought to be responsible for normal, healthy aging^5^. Animal studies further implicate SST+ neurons in the prelimbic cortex (PLC), a region highly involved in mood disorders and addiction, for their resiliency-conferring role in these disorders^6^. Interestingly, acute versus chronic inhibition of SST+ neurons in the PLC leads to opposing effects on behavioral emotionality^7^. These findings highlight the important yet complex functionality of these neurons. Unfortunately, whereas the human literature has placed a priority on detection of SST itself within CSF, the animal literature has focused largely on the GABAergic properties of SST+ neurons, with little focus on their peptidergic actions itself. Very little research has been conducted to elucidate circumstances of SST release, possibly due to difficulties in detection protocols. Here, we demonstrate a method for the detection of SST using commercially-available kits (and for the first time, experimentally induce both release and inhibition of SST) in artificial CSF (aCSF) of acute brain slices containing the PLC. Using a combination of optogenetics and enzyme-linked immunosorbent assay (ELISA)^8^ with transgenic mice, we optically induced SST release from SST+ neurons via activation of channelrhodopsin (ChR2) using physiologically relevant stimulation frequencies^9^. In addition, we show that inhibition of SST neurons via halorhodopsin (NpHR) reduces SST. This methodology can be adapted and translated to other SST+ neuronal populations based on their firing patterns, and applied across a wide variety of disease models. Our application provides a novel approach for future investigation into SST, and allows for greater translation between animal and clinical investigations of a multitude of psychiatric disorders.

## RESULTS

### Changes in SST release following optogenetic activation and inhibition (Fig. 1)

SST release and inhibition were reliably detected in brain slices. SST-IRES-Cre mice were injected with ChR2, NpHR, or mCherry in the PLC and given 3 weeks to allow for ample local expression of the virus. Injections were confined to the PLC (Fig. 1 B-C). Patch clamp experiments were conducted to confirm optically-evoked GABA currents (Fig. 1D). A 5 msec stimulation with a 470 nm laser induced GABA currents of approximately 80 pA. ChR2-induced SST release was detected following 10 Hz stimulation (one-way ANOVA, *F*_36,2_ = 5.164, *p* = 0.0107) (Fig. 1E**, left**). Tukey’s posthoc test revealed that 10 Hz stimulation led to a significantly increased detectable SST over control conditions (*p* = 0.0154) as well as over 5 Hz stimulation (*p* = 0.0315). 5 Hz stimulation did not differ from control (*p* = 0.9988). Because SST release was detected in mCherry control slices, suggesting tonic release of SST, we further demonstrated a 15 min continuous stimulation of NpHR led to significantly reduced SST release (paired *t*-test, *t*_7_ = 3.735, *p* = 0.0073) (Fig. 1E**, right**).

**Fig. 1:**
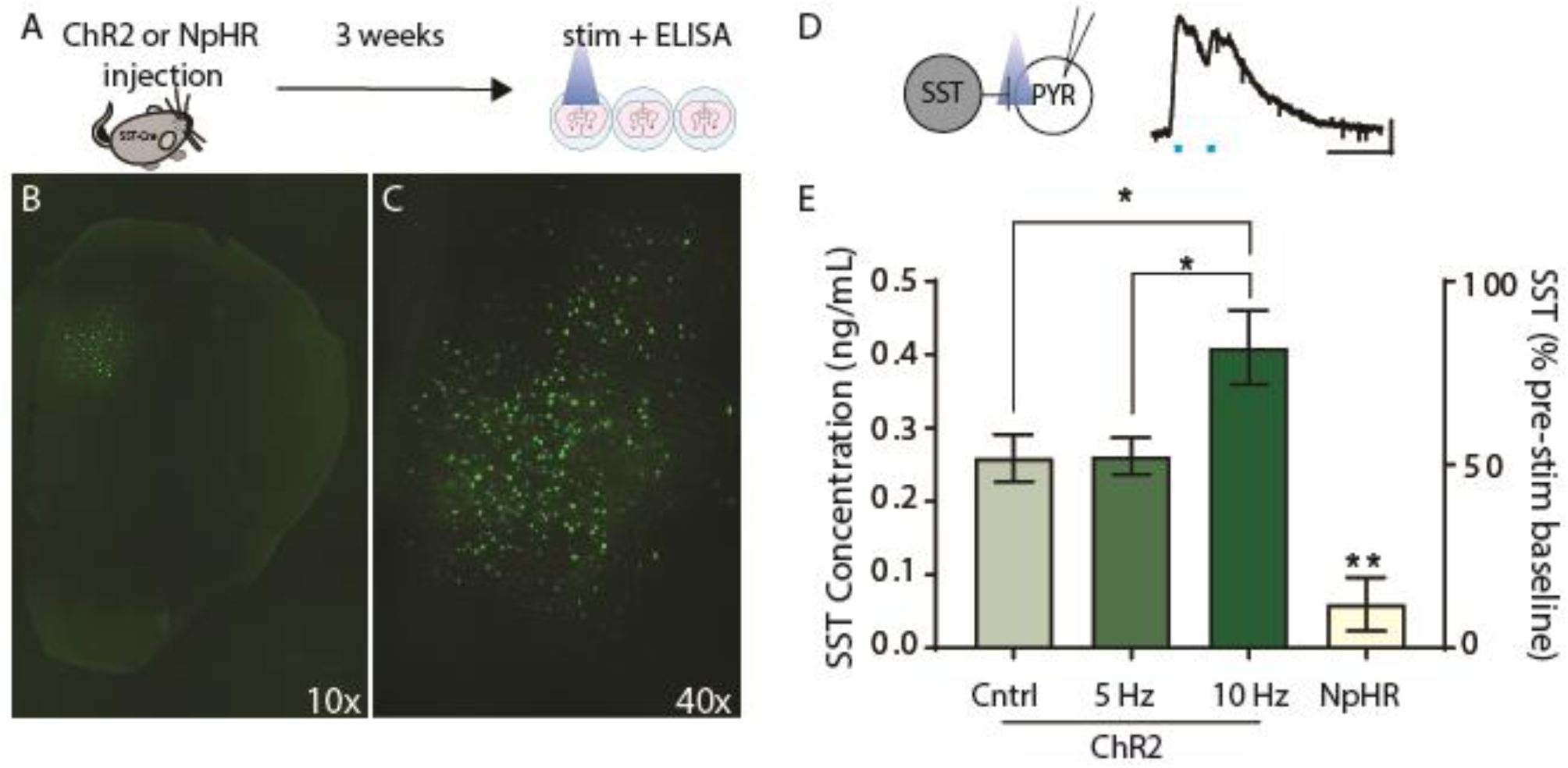
SST concentration is altered following optogenetic activation or inhibition of SST neurons. (A) Experimental design. (B-C) Representative images of ChR2 injections. ChR2 injections were confined to the prelimbic cortex (B-C). Representative traces confirming light-evoked GABA currents onto pyramidal neurons (D, scale bar represents 20 pA x 100 msec). Concentrations of SST in aCSF were increased following 10 Hz stimulation of ChR2 for 15 min (E). Inhibition via NpHR lead to a reduction in aCSF SST as compared to baseline.

## DISCUSSION

These experiments present a novel SST release protocol that is applicable to research programs across a broad array of neuropsychiatric disorders. Previous human literature has focused primarily on the detection of SST in CSF ^(1-2)^ and post-mortem brain tissue^10^. Within the preclinical animal literature, much of the work has focused on the overall neuronal activity and circuitry of SST+ neurons in affective regulation with little work on the SST peptide itself. Together, the SST+ neurons and SST peptide have emerged as critical cellular and molecular targets of neuropsychiatric disorders. However, these investigations have yet to elucidate the modulatory role of SST in neurotransmission within neuropsychiatric disease models. Additionally, the conditions in which GABA, potentially glutamate^11^, and SST are released from neurons remain elusive. Here we demonstrated that SST release from cortical SST+ neurons is enhanced during high frequency stimulation (10 Hz), concordant with previously known mechanisms of neuropeptide release^13^. Interestingly our data suggest that SST is also present under basal or tonic conditions, indicating a role for this peptide in regulating healthy brain function.

Our work combines optogenetics, transgenic mice, and ELISA to create a protocol for the induction of SST release and its detection in acute brain slices. We believe that the methodology presented here is easily adaptable for other brain regions enriched with SST. In addition, it can be applied to the variety of rodent disease models, such as those for depression (social defeat stress, unpredictable chronic mild stress, etc.). Future work should investigate whether this SST tone is necessary for healthy aging or the prevention of neuropsychiatric disorders such as MDD.

## METHODS

### Animals

6-week old SST-IRES-Cre male and female mice N=7, C57BL/6J genetic background, #013044, Jackson Laboratory) were bred in house. Mice were single-housed and maintained on a 12:12 h reverse light-dark cycle with food and water *ad libitum*. All procedures were approved by the Pennsylvania State University’s Institutional Animal Care and Use Committee (IACUC).

### Surgeries

Following craniotomy, mice were bilaterally injected using a 1µl Neuros syringe (65458-01, Hamilton Company, Reno, NV; stereotax, Stoelting) with 0.3 µl of the viral vectors into the PLC (AP: +1.80, ML: ±0.40, DV: −2.50). Bupivicaine was applied topically and ketoprofen intraperitoneally for postoperative analgesia. Mice were allowed to recover for three weeks prior to experiments for optimal expression.

### Enzyme Immunoassay for Photostimulated Release of SST

Acute brain slices for photostimulation were prepared as previously described^8^. Briefly, mice were rapidly decapitated under isofluorance anesthesia and the dissected brain was placed in ice-cold modified high-sucrose artificial cerebrospinal fluid^12^. 150 µM slices containing the PLC were dissected on a Compresstome (Precisionary Instruments, Greenville, NC) and transferred to normal aCSF for 30 min. Slices were maintained in a holding chamber at 31°C, continuously bubbled with a 95% O_2_/ 5% CO_2_ mixture and shielded from light until photostimulation. Individual slices were placed in a well on a 12-well plate with 500 uL of normal aCSF. For the ChR2 optical stimulation experiment, a 470-nm laser (CoolLED, United Kingdom) was directed to the slice for 15 min (5 Hz and 10 Hz). Stimulation frequency was counterbalanced between rostral and caudal PLC slices to account for variability in viral expression along the anterior/posterior axis. At the end of the experiment, 3×50 uL of the aCSF within the well was pipetted into a 96-well SST ELISA plate (Peninsula Labs, cat. #S-1179) as per the manufacturer’s instruction. Absorbance was read at 450 nm on a Synergy2 microplate reader (BioTek). SST release was quantified in ng/mL. A similar procedure was done for the NpHR optical inhibition experiment, except that a baseline SST concentration level was taken and new aCSF placed in the well plate. After this, a 565-nm laser was directed to the well continuously for 15 min.

### Viral Vectors

The viral constructs AAV5-hSyn-DIO-mCherry (#50459), AAV5-EF1a-DIO-hChR2(H134R)-eYFP-WPRE-HGHpA (#20298), and AAV5-Ef1a-DIO-eNpHR3.0-eYFP (#26966) were obtained from Addgene (Watertown, MA).

### Electrophysiology

Electrophysiology experiments were conducted as published^12^. 300 µM coronal slices containing the PLC were prepared as described above. A cesium-methanesulfonate based intracellular recording solution was used to record light-evoked inhibitory events were at +10mV.

### Histology

Following photostimulation, slices were immersed in 4% paraformaldehyde overnight, mounted on slides and coverslipped. Images were obtained on an epifluorescent microscope to verify viral expression of EYFP in the PLC.

### Statistics

All data are represented as mean ± SEM. Statistical analyses were performed in Graphpad Prism 7.0. A one-way ANOVA was used for ChR2 experiments, followed by Tukey’s posthoc test. NpHR experiment was analyzed by paired Student’s t-test. Statistical significance was taken as \**p* < 0.05.

## ACKNOWLEDGEMENTS

This work was funded by The Brain and Behavior Research Foundation NARSAD Young Investigator Award (NAC), and Penn State University’s Social Science Research Institute (NAC). We would like to acknowledge Dr. Janine Kwapis (Penn State University) for her expertise in imaging.

## References

1. Hannon JP, Nunn C, Stolz B, et al. Drug Design at Peptide Receptors. J Mol Neurosci. 2002;18(1-2):15–28. doi:10.1385/JMN:18:1-2:15

2. Ge W, Zhou D, Zhu L, Song W, Wang W. Efficacy and Safety of Everolimus plus Somatostatin Analogues in Patients with Neuroendocrine Tumors. J Cancer. 2018;9(24):4783–4790. doi:10.7150/jca.25908

3. Agren H, Lundqvist G. Low levels of somatostatin in human CSF mark depressive episodes. Psychoneuroendocrinology. 1984;9(3):233–248. http://www.ncbi.nlm.nih.gov/pubmed/6149588. Accessed March 21, 2019.

4. Bissette G, Widerlöv E, Walléus H, et al. Alterations in cerebrospinal fluid concentrations of somatostatinlike immunoreactivity in neuropsychiatric disorders. Arch Gen Psychiatry. 1986;43(12):1148–1151. http://www.ncbi.nlm.nih.gov/pubmed/3778111. Accessed March 21, 2019.

5. Sibille E. Molecular aging of the brain, neuroplasticity, and vulnerability to depression and other brain-related disorders. Dialogues Clin Neurosci. 2013;15(1):53–65. http://www.ncbi.nlm.nih.gov/pubmed/23576889. Accessed March 21, 2019.

6. Fuchs T, Jefferson SJ, Hooper A, Yee P-H, Maguire J, Luscher B. Disinhibition of somatostatin-positive GABAergic interneurons results in an anxiolytic and antidepressant-like brain state. Mol Psychiatry. 2017;22(6):920–930. doi:10.1038/mp.2016.188

7. Soumier A, Sibille E. Opposing effects of acute versus chronic blockade of frontal cortex somatostatin-positive inhibitory neurons on behavioral emotionality in mice. Neuropsychopharmacology. 2014;39(9):2252–2262. doi:10.1038/npp.2014.76

8. Al-Hasani R, McCall JG, Shin G, et al. Distinct Subpopulations of Nucleus Accumbens Dynorphin Neurons Drive Aversion and Reward. Neuron. 2015;87(5):1063–1077. doi:10.1016/j.neuron.2015.08.019

9. Yavorska I, Wehr M. Somatostatin-Expressing Inhibitory Interneurons in Cortical Circuits. Front Neural Circuits. 2016;10:76. doi:10.3389/fncir.2016.00076

10. Tripp A, Kota RS, Lewis DA, Sibille E. Reduced somatostatin in subgenual anterior cingulate cortex in major depression. Neurobiol Dis. 2011;42(1):116–124. doi:10.1016/j.nbd.2011.01.014

11. Cattaneo S, Zaghi M, Maddalena R, Bedogni F, Sessa A, Taverna S. Somatostatin-Expressing Interneurons Co-Release GABA and Glutamate onto Different Postsynaptic Targets in the Striatum. bioRxiv. March 2019:566984. doi:10.1101/566984

12. Crowley NA, Bloodgood DW, Hardaway JA, et al. Dynorphin Controls the Gain of an Amygdalar Anxiety Circuit. Cell Rep. 2016;14(12):2774–2783. doi:10.1016/j.celrep.2016.02.069

13. Crowley NA, Bloodgood DW, Hardaway JA, et al. Dynorphin Controls the Gain of an Amygdalar Anxiety Circuit. Cell Rep. 2016;14(12):2774–2783. doi:10.1016/j.celrep.2016.02.069

